# Optimization and deoptimization of codons in SARS-CoV-2 and the implications for vaccine development

**DOI:** 10.1101/2022.09.03.506470

**Authors:** Xinkai Wu, Kejia Shan, Fuwen Zan, Xiaolu Tang, Zhaohui Qian, Jian Lu

## Abstract

The spread of Coronavirus Disease 2019 (COVID-19), caused by the SARS-CoV-2 coronavirus, has progressed into a global pandemic. To date, thousands of genetic variants have been identified across SARS-CoV-2 isolates from patients. Sequence analysis reveals that the codon usage of viral sequences decreased over time but fluctuated from time to time. In this study, through evolution modeling, we found that this phenomenon might result from the virus’ preference for mutations during transmission. Using dual luciferase assays, we further discovered that the deoptimization of codons on viruses might weaken protein expression during the virus evolution, indicating that the choice of codon usage might play important role in virus fitness. Finally, given the importance of codon usage in protein expression and particularly for mRNA vaccine, we designed several omicron BA.2.12.1 and BA.4/5 spike mRNA vaccine candidates based on codon optimization, and experimentally validated their high levels of expression. Our study highlights the importance of codon usage in virus evolution and mRNA vaccine development.

## Introduction

Amino acids are the building components of proteins. Eighteen out of the twenty amino acids are encoded by at least two synonymous codons in all domains of life, with the exception of methionine and tryptophan (*1, 2*). The synonymous codons of an amino acid are frequently used at different frequencies because of the host cell’s variable tRNA supply, which is referred to as “codon usage bias” (CUB). Most organisms have some degree of CUB (*3*). Due to the degeneracy of the genetic code, synonymous mutations in protein-coding regions are frequently considered neutral or almost neutral since they do not alter the protein sequence (*4, 5*). However, accumulating evidence suggests that synonymous mutations are not entirely neutral in evolution (*1, 6-8*). By altering translation efficiency (*9*), mRNA stability (*10*), and peptide conformation (*11, 12*), synonymous mutations may impact the protein expression and function, and eventually the fitness of organisms (*6*).

Viruses typically rely on the cellular machinery of the host organisms for biological processes such as translation. Viruses usually exhibit a modest level of CUB, presumably due to mutational pressure (*13, 14*). It is likely that viruses with low and inefficient codon usage can adapt to various host species with different codon usage preferences (*14-16*). The host cellular environment frequently attacks the RNA virus genomes, which can accumulate specific types of mutations (*17*). For instance, the APOBEC family can produce C>U mutations in the viral RNA genomes, while the ADAR family can result in A>G mutations (*18-20*). It was demonstrated that codon usage patterns in RNA viruses might be primarily influenced by mutational pressure (*14, 16, 21-25*). Natural selection may also play an important role in modulating viral CUB, although its effectiveness may vary amongst viruses. For instance, natural selection has significantly influenced how codons are used in coronaviruses (*26*). Furthermore, too much CUB similarity between viruses and hosts will negatively impact both the viruses and the hosts (*7*).

SARS-CoV-2, the etiology of the Coronavirus Disease 2019 (COVID-19), has a codon usage pattern different from that of humans and bats (*15, 27-30*). Studies have revealed that as the pandemic spread, SARS-CoV-2’s codon usage pattern diverged further from that of its human hosts rather than evolving to use more optimized codons (*31, 32*). Despite these discoveries, the evolutionary driving forces underlying the optimization and deoptimization of codons in SARS-CoV-2 are poorly understood. Although some studies suggest that natural selection is the main force shaping the codon use patterns in SARS-CoV-2, others argue that mutational pressure is the primary factor determining the codon usage of this virus and natural selection only played a minor role (*33*). Additionally, some studies suggest that the codon usage of SARS-CoV-2 is influenced by both mutational bias and natural selection (*28, 29*). Therefore, significant gaps remain regarding the evolutionary principles of CUB in SARS-CoV-2. Furthermore, experimental evidence that links codon usage to the translational efficiency of SARS-CoV-2 genes is still lacking.

Here, we analyzed 9,164,789 high-quality SARS-CoV-2 genomes for codon usage profiles. We found the deoptimization of codons in the evolution of SARS-CoV-2 genomes is primarily influenced by the amino acid changes, while synonymous mutations had little effect. We identified a strong bias toward C>U substitutions, and some of them might be favored by natural selection. We showed experimentally that codon optimization of SARS-CoV-2 improves protein translation in human cells. Finally, we showed that optimizing the Spike gene codon usage of SARS-CoV-2 variants has important implications for vaccine design.

## Results

### The deoptimization of codons is mainly caused by amino acid changes in the continuing evolution of SARS-CoV-2

We downloaded 9,164,789 high-quality SARS-CoV-2 genomes from the Global Initiative on Sharing All Influenza Data (GISAID, https://www.gisaid.org; as of June 18, 2022), and calculated the Codon Adaptation Index (CAI) of the concatenated coding sequences of each genome. The CAI value ranged from 0.6154 to 0.6192, with a median value of 0.6166 and the 2.5th and 97.5th percentiles of 0.6162 and 0.6169, respectively. To decipher the evolutionary trend of CAI as the pandemic developed, we compared the CAI values of the SARS-CoV-2 genomes collected on different dates. Consistent with previous findings that CAI decreased during the first four (*31*) and 18 months (*32*) of SARS-CoV-2 evolution in humans, we also observed such a pattern (**Figure 1a**, the red lines). However, we observed a turning point of CAI around November 26, 2021, roughly 23 months after the initiation of the COVID-19 pandemic. Before the turning timepoint, the CAI value decreased as the Alpha variants (B.1.1.7) became the predominant lineage, and the CAI value further dropped as the Delta (B.1.617.2) variants replaced the Alpha and other variants of concern (VOCs) or variants of interest (VOIs). After November 26, 2021, the Omicron variants (B.1.1.529) rapidly replaced the Delta variants and became the predominant SARS-CoV-2 variant, and correspondingly, the CAI value of SARS-CoV-2 increased (**Figure 1a, b**).

**Figure 1.**
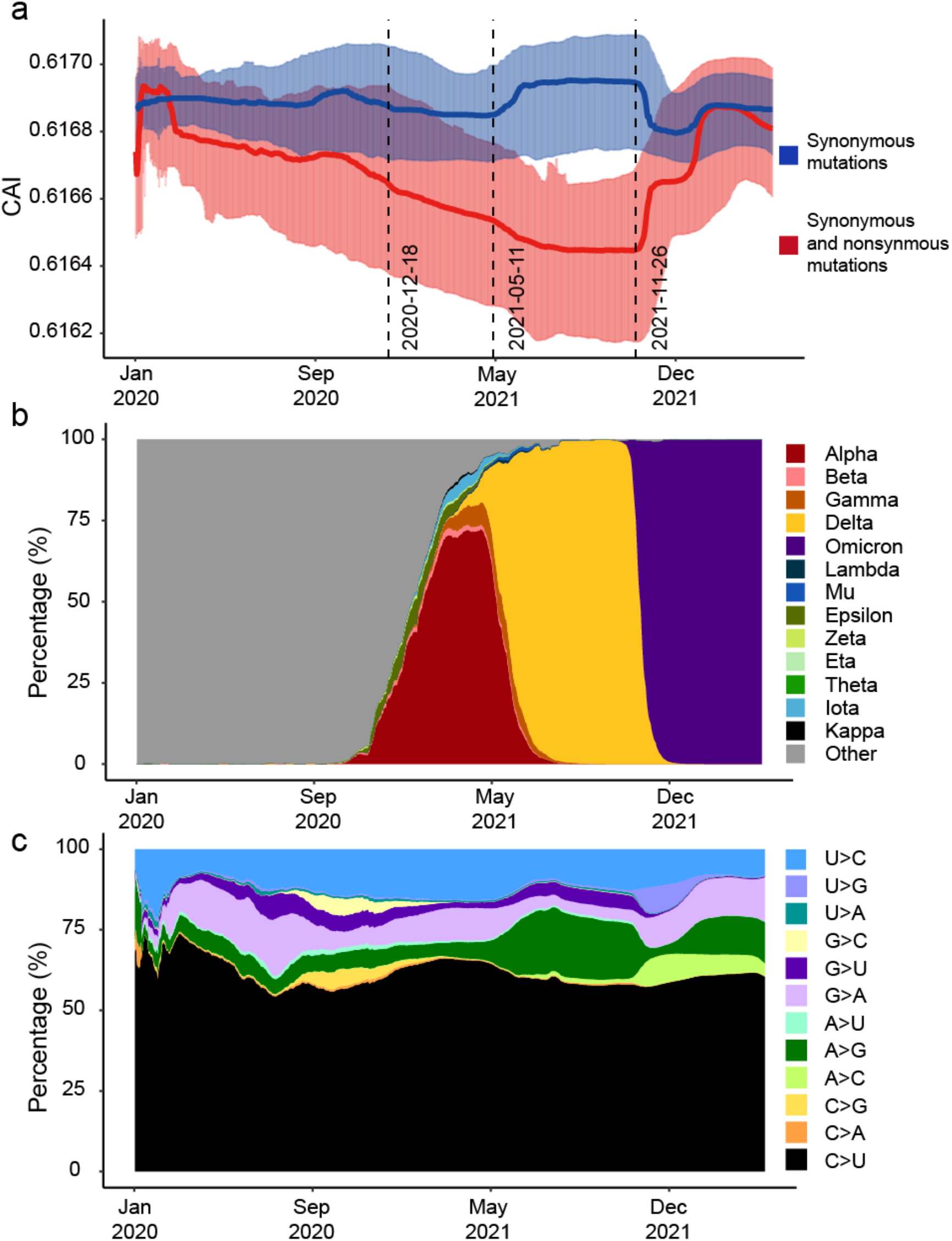
The evolutionary trends of CAI, major viral lineages, and synonymous mutational types during SARS-CoV-2 evolution. (a) The CAI change of SARS-CoV-2 caused by synonymous and nonsynonymous (the solid red line and shadow) or synonymous mutations (the solid blue line and shadow) over time. The red and blue solid lines indicate the median of CAI, and the red and blue shadows indicate the 95% interval of CAI in a 14-days window with a one-day step. The black dash lines indicate the dates when World Health Organization defined the Alpha, Delta, and Omicron lineages as the variants of concern. (b) Prevalence of VOCs and VOIs over time. The proportion of variants of SARS-CoV-2 in a 14-days window with a one-day step. (c) The proportions of synonymous mutations of different substitutional types in SARS-CoV-2 in a 14-days window with a one-day step.

As the pandemic progressed, positive selection has driven many amino acid substitutions in the SARS-CoV-2 genomes (*34*). In particular, the Alpha, Delta, and Omicron variants differ from the reference genome (NC_045512) by at least 17, 26, and 31 amino acids (retrieved from Outbreak.info project, https://outbreak.info/), respectively. To examine whether the decreasing or increasing of CAI values was attributable to the amino acid changes in the VOC/VOI variants, we neglected the nonsynonymous changes that alter amino acids and calculated the CAI value of each genome (only the synonymous changes were considered in the CAI analysis). Interestingly, we found that the CAI value fluctuates slightly in a tight range between 0.6168 and 0.6170 (**Figure 1a**, blue lines). In other words, synonymous changes only slightly affected the CAI values of SARS-CoV-2 throughout the pandemic, and the relatively large decreases or increases in CAI mainly resulted from nonsynonymous substitutions in SARS-CoV-2. We hypothesize that the overall change in CAI value among SARS-CoV-2 variants may result from coincidental by-products of amino acid changes that may affect the replication, transmission, or immune evasion of SARS-CoV-2.

### Deoptimization of codons by C>U substitutional bias in SARS-CoV-2

The APOBEC RNA editing enzymes frequently induce C>U mutations in the single-stranded viral RNA molecules (*35-40*), causing excessive C>U substitutions in the SARS-CoV-2 genomes (*38, 41-45*). Notably, the C>U substitutions, which roughly accounted for ∼60% of all the nucleotide changes accumulated in the synonymous sites in the SARS-CoV-2 genomes, remained dominant over time despite the frequent replacement of viral lineages (**Figure 1c**).

To investigate the possible impact of C>U synonymous mutations on CAI, we computed the *w*_*ij*_ parameter to measure the relative usage of a codon (*j*) among all the codons that encode a specific amino acid (*i*). The *w*_*ij*_ parameter was calculated based on the occurrences of codons in the coding sequences and the gene expression profiles in 54 human tissues (Materials and Methods). Thus, *w*_*ij*_ = 1 means codon *j* is the most used in the transcriptomes among all the codons encoding amino acid *i*. Notably, all the synonymous changes generated by C>U deoptimize SARS-CoV-2 codons (that is, to reduce the *w* value). The top 30 most abundant synonymous codon changes accounted for 90.7% of all the synonymous codon changes that differ by one nucleotide in the analyzed SARS-CoV-2 genomes, and 16 (53.3%) of such codons were caused by C>U mutations (**Figure 2a**). Consequently, the C>U substitutional bias considerably contributed to the decreased CAI value in SARS-CoV-2.

**Figure 2.**
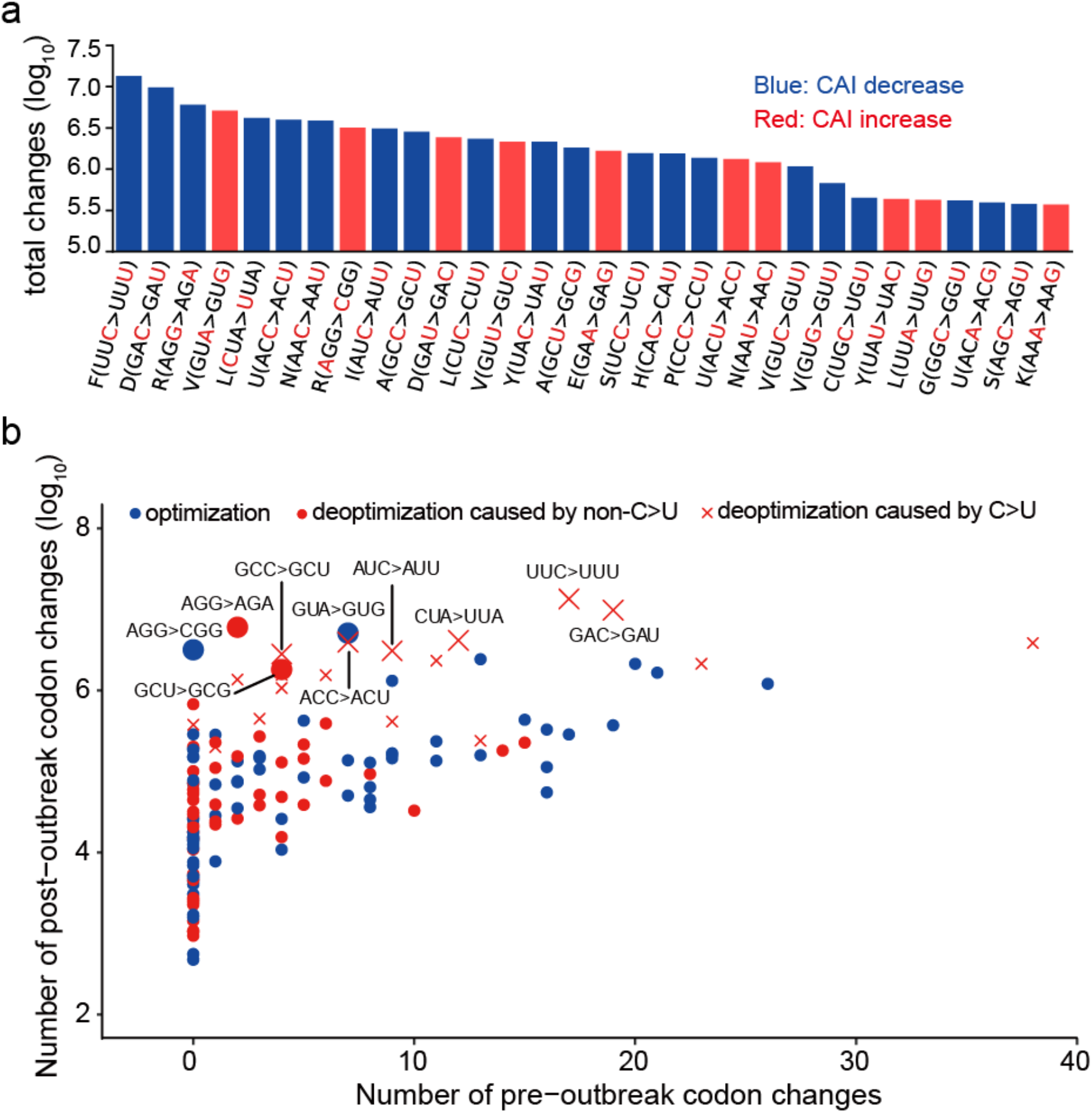
Deoptimization of codons by C>U substitutional bias in SARS-CoV-2. (a) Type of the top 30 observed codon substitutions by frequency. Red denotes an increase in the sequence’s CAI, whereas blue suggests a loss as a result of the substitution. (b) The number of pre- or post-outbreak synonymous codon changes. The *x*-axis indicates the number of pre-outbreak codon changes from the ancestor to SARS-CoV-2 and RaTG13. The *y*-axis indicates the number of codon changes that occurred during the pandemic. A blue dot indicates an unpreferred codon was replaced by a preferred one (optimization). A red dot indicates apreferred codon was replaced by an unpreferred one (deoptimization). The red x shapes indicated the codon deoptimization caused by C>U mutations. The ten synonymous codon changes occurred more frequently during the pandemic than in pre-outbreak history (Fisher’s exact test, *P*-adj < 0.05) were labeled, and the relative dots or x shapes were enlarged.

Nucleotide substitutional bias was observed in the evolution of coronaviruses closely related to SARS-CoV-2 prior to the COVID-19 pandemic (*18-20, 46*). To examine whether the relative frequencies of nucleotide substitutional types differ between the pre-outbreak history in animals and the continuing evolution of SARS-CoV-2 in humans, we inferred the ancestral state of each codon in the most recent common ancestor of RaTG13 and SARS-CoV-2 as previously described (*46*), and counted the frequencies of synonymous codon changes from the ancestral state to RaTG13 or SARS-CoV-2 (to reduce the stochastic noise, we pooled the synonymous changes in the two branches). As shown in **Figure 2b**, the occurrences of synonymous codon changes were significantly positively correlated in the pre-outbreak evolution history in animals and the post-outbreak evolution of SARS-CoV-2 in humans (Spearman’s *ρ* = 0.73, *P* < 10^−10^). We found at least ten synonymous codon changes significantly enriched in SARS-CoV-2 than in pre-outbreak SARS-CoV-2 like viruses, and notably, 6 of them are C>U changes that tend to deoptimize codon usage (**Figure 2b**). Among the four remaining types of synonymous codon change, one is G>A substitution, which might be caused by C>U editing in the antisense; and one is the A>G change, which is putatively caused by the A-to-I editing mediated by ADAR enzymes (**Figure 2b**). Considering the genome sequence of BANAL-20-52 recently isolated from Laos is most similar to SARS-CoV-2 (*47*), we also obtained the synonymous codon changes from the most recent common ancestor of BANAL-20-52 and SARS-CoV-2 to them in the pre-outbreak evolution history (**Figure S1**). The excessive synonymous mutations due to C>U substitution were still observed in the SARS-CoV-2 genomes, although two C>U changes (GCC>GCU and AUC>AUU) were not enriched in the SARS-CoV-2 genomes (**Figure S1**). Overall, these results suggest that excessive C>U changes have accumulated in the SARS-CoV-2 genomes after shifting the host from animals to humans, presumably as a result of different host environments.

### The high-frequency synonymous substitutions in SARS-CoV-2 are dominated by C>U changes

To explore the evolutionary forces acting on the C>U synonymous changes, we first calculated the derived allele frequency (DAF) of the synonymous changes in all the SARS-CoV-2 genomes. In total, 19,004 distinct synonymous changes in 10,214 sites were observed, with 5,473 sites having one type of synonymous change, 692 sites showing two types of synonymous changes, and 4,049 sites exhibiting three types of synonymous changes, indicating that many of these synonymous changes are the result of independent recurrent mutations. The DAF ranged from 1.09×10^−7^ to 0.994, with a median value of 2.22×10^−5^ and the 2.5% and 97.5% quantiles of 2.18×10^−7^ and 1.40×10^−3^, respectively. Notably, the C>U synonymous changes had significantly higher DAFs than the non-C>U synonymous changes (*P* < 10^−16^), and the median DAF of the former was more than 20 times that of the latter (3.75×10^−4^ versus 1.80×10^−5^, **Figure 3a**). We also utilized a second method to minimize the potential bias in estimating DAFs. Briefly, we binned the SARS-CoV-2 genomes collected every 14 days, calculated the DAFs of the synonymous changes in each bin, and used the maximum DAF of a specific change to represent its DAF. Using this method, we still found the DAFs of the C>U synonymous changes are significantly higher than those of the non-C>U synonymous changes (*P* < 10^−16^), and that the median DAF of the former was 13.1 times that of the latter (1.84×10^−3^ versus 1.41×10^−4^, **Figure 3b**). These patterns persisted when we separately analyzed the Delta or Omicron variants (**Figure S2**). These findings suggest that the C>U synonymous substitutions, which typically deoptimize the codons, have higher allele frequencies in the SARS-CoV-2 populations than the non-C>U synonymous substitutions.

**Figure 3.**
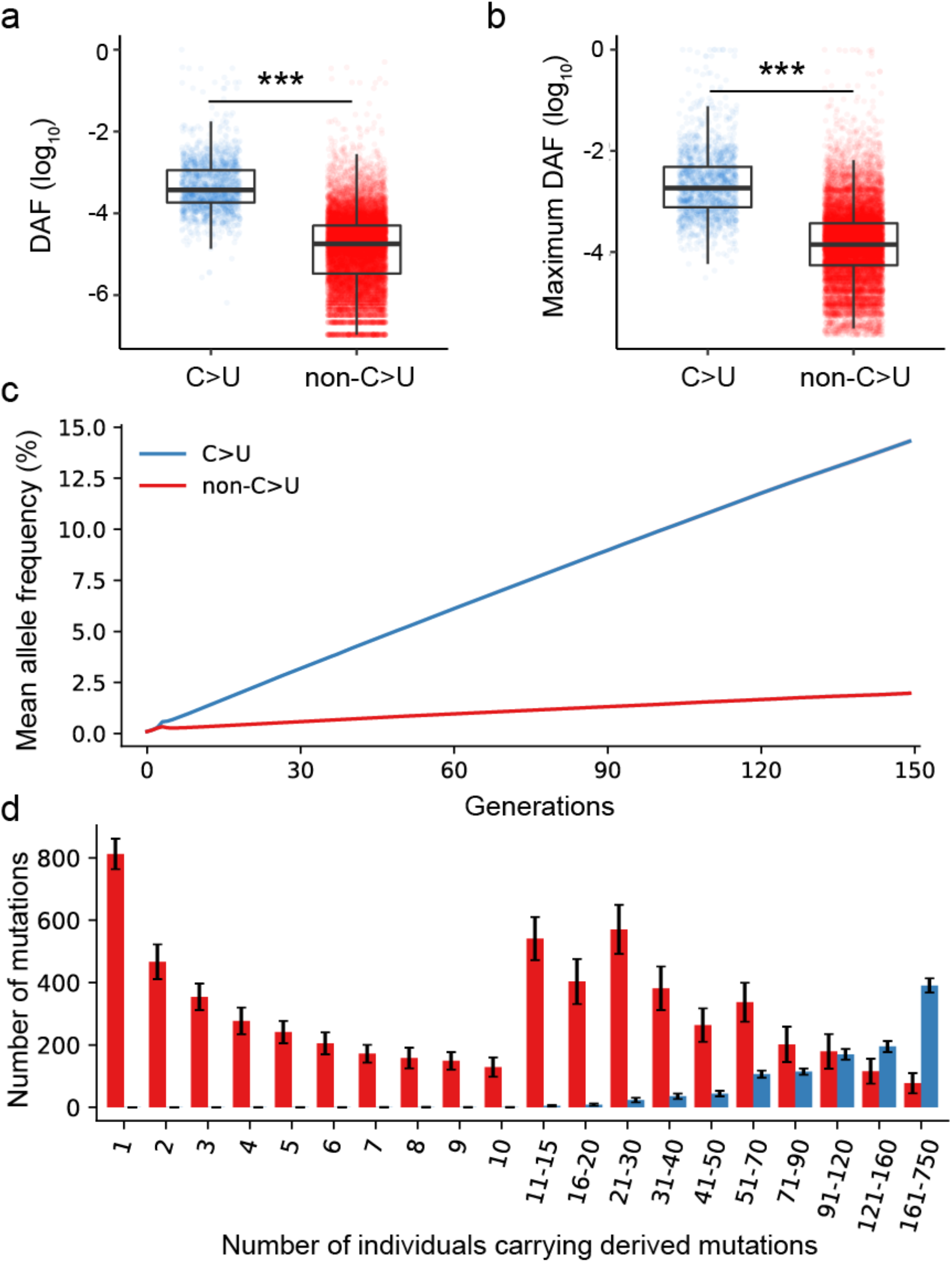
Simulation of viral sequences under biased mutation and random selection. (a) The C>U synonymous mutations had significantly higher DAFs than the non-C>U synonymous changes. (b) The C>U synonymous changes had significantly higher maximum DAF than the non-C>U synonymous changes. (c) Allele frequency of C>U and non-C>U mutations in the population during the simulation. The *x*-axis is generations in the simulation process, and the *y*-axis is the frequency (mean) of the C>U or non-C>U mutations observed in the population. (d) The allele frequency spectrum of C>U and non-C>U mutations in the 150thgeneration. Mutations were grouped rationally according to their observed number in the population, and the number of mutations in each group was counted. The average number of mutations in each group over 100 independent simulations was calculated and plotted in the graph.

To explore whether mutational bias alone can cause elevated DAFs of C>U synonymous mutations, we conducted simulations with a biased sequence evolution model (Materials and Methods). In brief, our simulation considered the mutational bias based on the number of synonymous substitutions accumulated in the SARS-CoV-2 populations (**Table 1**). In the simulations, neither natural selection nor recombination was taken into account, and the whole sequence evolution process was replicated 100 times. As shown in **Figure 3c**, although the mean DAF increases for both C>U and non-C>U mutations as the number of generations increases, the mean DAF tends to be higher for C>U mutations, presumably resulting from independent recurrent mutations on the same sites. Based on the mutation spectra obtained in our simulations, C>U mutations tend to be enriched in the higher frequency category (the frequency spectra of mutations at generation 150 are shown in **Figure 3d**). Nevertheless, the DAF did not differ more than ten times between these two categories of mutations in our simulations, as observed in actual SARS-CoV-2 populations (**Figure S3**). For example, at the 150th generation in our simulations, the mean DAF was 14.3% for C>U mutations, while the corresponding number was 2.0% for the non-C>U mutations. Thus, although C>U mutation bias can lead to high frequencies for C>U mutations, natural selection might need to be invoked to account for the observed difference in DAF between the C>U and other types of synonymous changes.

The role of natural selection on C>U synonymous mutations is yet unclear (*44, 45*). The DAFs of synonymous changes are affected by either sampling bias in SARS-CoV-2 genome sequencing, or by the frequent replacement of the SARS-CoV-2 lineages over the pre-existing ones during the COVID-19 pandemic. Consequently, it is challenging to estimate the distribution of synonymous mutational effects in the absence of strictly neutral evolving sites. Nevertheless, it will be unsurprising that many of the C>U and non-C>U synonymous mutations are subject to very strong purifying selection, given the extremely low DAF values of these mutations. On the other hand, we found 29 synonymous mutations that have DAF values greater than the mean + 3×s.d. (0.0343) of the total DAF values (**Table S1**), with C>U significantly over-represented in these changes than non-C>U changes (18/1670 versus 11/17334; *P* = 1.4×10^−12^, Fisher’s exact test). It is likely these high-frequency synonymous changes have been driven by positive selection.

### Experimental verifying translational regulation of synonymous changes

The synonymous mutations potentially influence protein translation efficiency by optimizing or deoptimizing the codons. To determine whether synonymous mutations in SARS-CoV-2 affect protein expression, we chose 34 synonymous mutations at random and used a dual-luciferase reporter assay (psiCHECK-2 vector) to analyze their effects. Briefly, for each synonymous mutation, we made two reporter plasmids with the wildtype (WT) or mutant (MUT) allele and the flanking sequences (120 nucleotides on each flank) of SARS-CoV-2 right after the start codon of Renilla **(Figure 4a**). The construction of the plasmid could be divided into two main steps: 1) linearization of the backbone DNA, and 2) synthesis of the insert fragment. By designing primers upstream and downstream of the insertion site for reverse PCR, we obtained the entire linearized plasmid backbone. We designed primers by segmentation and successfully constructed the insert fragment (including WT and MUT) sequences by overlap PCR. Finally, we assembled the insert and linearized backbone by Gibson assembly and successfully obtained WT and MUT vectors (Materials and Methods). The expression of Renilla in the psiCHECK-2 would be altered if the mutation had an effect on protein synthesis. We measured the fluorescence intensities of Renilla and Firefly. By contrasting Renilla’s expression levels between the WT and MUT vectors using Firefly as the reference, we calculated the effect of the occurrence of mutations on protein expression.

**Figure 4.**
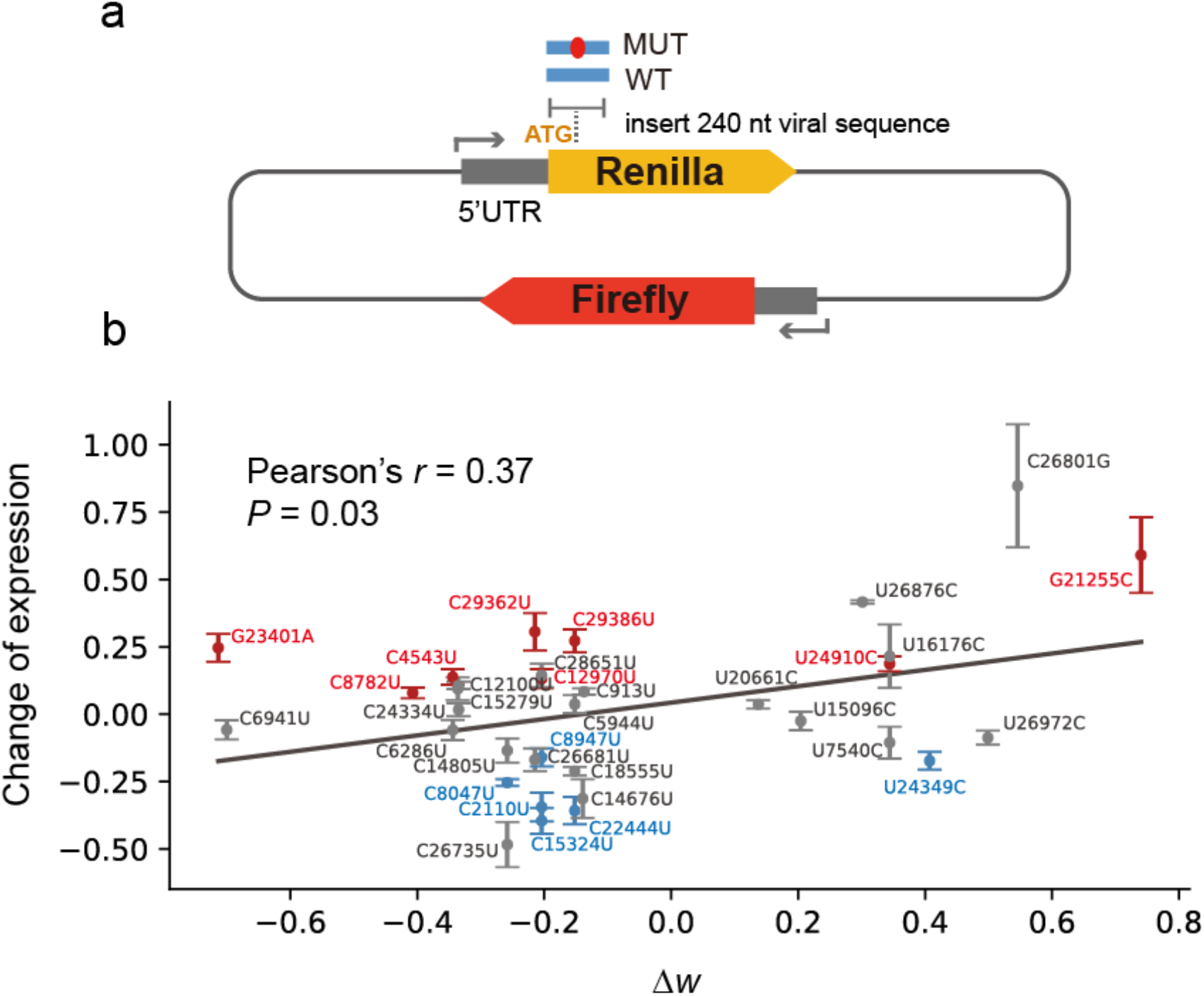
Effects of synonymous viral mutations on protein synthesis rate. (a) The design of dual luciferase reporter assays. By inserting a 240 nt viral CDS sequence (centered on the mutation) after Renilla’s start codon, two reporter plasmids (WT and MUT) were created. The function of the mutation was inferred by comparing the protein expression differences between MUT and WT. (b) The correlations between protein expression changes and codon usage change by synonymous mutations. The *y*-axis is the log2 ratio of protein production of the mutant allele relative to the wildtype allele. The *x*-axis represents the change of *w* after codon substitution, with a total of 34 pairs of WT and MUT. The median and standard errors of the change in protein expression level are presented for each mutation. Points are marked in red or blue when the relative intensity of the mutant allele is significantly higher or lower than that of the wildtype allele (two-sided t-test, P-adj < 0.05), and in gray if there is no significant difference.

Among the 34 synonymous mutations we tested, 24 of them changed non-optimized (*w*_*ij*_ < 0.9) codons to optimized (*w*_*ij*_ > 0.9) ones, and the remaining ten mutations changed optimized codons to non-optimized ones. We discovered a significant positive correlation between the change of protein expression level and the change in *w* (Δ*w*) for a synonymous mutation (Pearson’s *r* = 0.37, *P* = 0.03; **Figure 4b**), supporting the notion that codon optimization tends to increase viral protein synthesis and codon deoptimization tends to decrease viral protein synthesis. We found that 8 of the 34 synonymous mutations significantly increase protein translation (red points in **Figure 4b**) and six significantly decrease protein translation (blue points in **Figure 4b**). Notably, we previously discovered that, based on two tightly mutations (the C8782U synonymous mutation and the U28144C nonsynonymous mutation), the SARS-CoV-2 genomes could be divided into S lineage (U8782 and C28144) and L lineage (C8782 and U28144), with the S lineage being ancestral and the L lineage being derived from the S lineage (*48*). According to our luciferase reporter assay, the derived C8782 allele found in the reference genome (L lineage) has lower translational efficiency than the ancestral U8782 allele found in the S lineage. Furthermore, the C15324U mutation that had a high frequency in the B.1 variant popular in Basel, Switzerland in early 2020, decreases protein expression levels. In addition, the C29362U mutation that had high frequency in the Epsilon variants increases protein expression.

### Optimizing codons of the *S* gene of SARS-CoV-2 for vaccine designing

Codon selection is an essential consideration in vaccine design because different codon usage affects the efficiency of protein production in vaccine development, affecting antigen expression(*49*). The Spike (S) protein of coronaviruses binds the host receptor to allow viral entry into host cells and determines host ranges and tissue tropism. Moreover, the S protein is also the major target for neutralizing antibodies produced by host organisms. Given the importance of the S protein, many mRNA and protein subunit vaccines of SARS-CoV-2 are designed to target it. However, the S coding sequences from SARS-CoV-2 genomes of the reference genome and the VOC/VOIs had low CAIs. For example, the CAI of the *S* gene was 0.624 for the reference genome, and 0.624 and 0.626 for the BA.2.12.1 (BA.2.12.1-s1) and BA.4/BA.5 (BA.4/5-s1) variants in the Omicron lineage, respectively. On the other hand, human genes had a median CAI value of 0.742, with 2.5th and 97.5th percentiles of 0.636 and 0.842, respectively **(Figure 5a**). These findings highlighted the importance of optimizing the S gene’s codon usage for vaccine design against SARS-CoV-2 in human or other mammalian cells.

**Figure 5.**
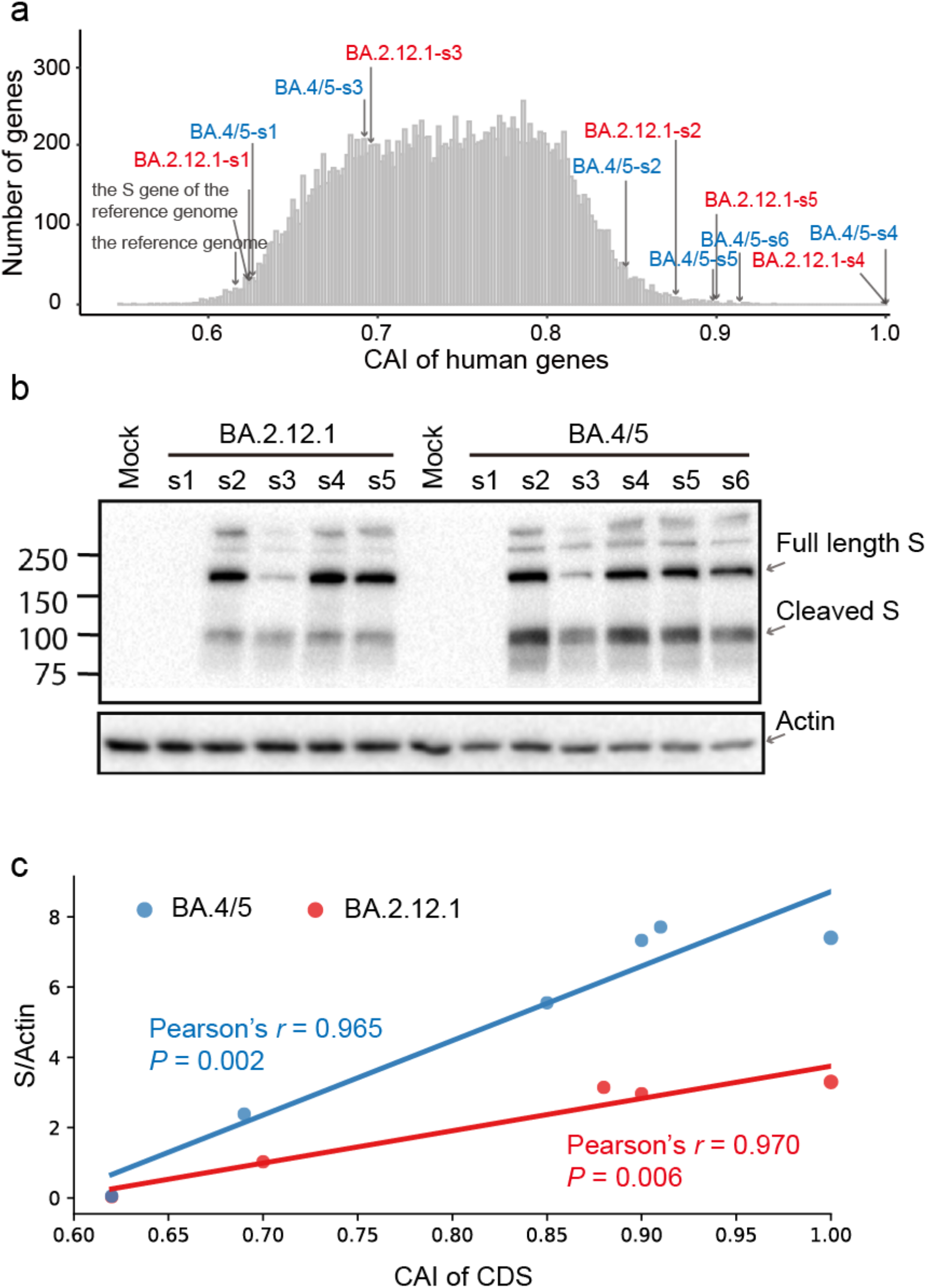
Experimental validation of codon optimization of the S gene. (a) The distribution of CAI values of human genes and the partially or fully optimized codons for the S gene of SARS-CoV-2. For the S gene of BA.2.12.1 and BA.4/5 variants in the Omicron lineage, s1 represents the native viral CDS sequence, s2, s3, and s5 represent the sequences with different CAI optimization levels, respectively, and s4 represents the fully optimized CDS sequence that had a CAI value of 1. (b) Western blotting analysis of S protein expression in HEK293T cells. The bands of full-length and cleaved S protein are labeled. (c) Correlation analysis between CAI value and S protein expression level. Red, BA.2.12.1; blue, BA.4/5.

To further evaluate the effect of codon usage on S protein expression, we developed an algorithm for codon optimization and constructed a number of plasmids encoding the S protein of the Omicron variants with various levels of optimized mammalian codon usage. We included S proteins of the BA.2.12.1 and BA.4/5 variants in our study. For each variant S protein, the encoding amino acid sequences from different constructs were identical, whereas their CAI values were different **(Figure 5a**). The levels of their expression were compared with the constructs with the native sequence derived from the viral genome (BA.2.12.1-s1 and BA.4/5-s1) and the fully-optimized sequence with a CAI value of 1 (BA.2.12.1-s4 and BA.4/5-s4) by western blot analysis. For each S protein, we also added three randomly selected partially-optimized sequences (s2, s3, and s5) with CAI values ranging from 0.68 to 0.95 in the analyses. As shown in **Figure 5b**, the viral original S protein coding sequence only showed a background level of S protein expression (BA.2.12.1-s1 in lane 2, and BA.4/5-s1 in lane 8). In contrast, the fully optimized sequences gave a high level of S protein expression for both the BA.2.12.1 (BA.2.12.1-s4 in lane 5) and BA.4/5 (BA.4/5-s4 in lane 11) variants. Overall, the levels of S protein expression were strongly correlated with the CAIs (**Figure 5c**). Thus, these results highlighted the significance of codon usage optimization in mRNA and DNA vaccine design.

## Discussion

Typically, viruses rely on the cellular machinery of their hosts for biological functions such as translation. The codon usage patterns utilized by the viruses could impact their protein production in host cells, ultimately affecting their replication and transmission. However, the functional consequences of synonymous mutations in SARS-CoV-2 are under-studied and poorly understood. In this work, we showed that the CAI of SARS-CoV-2 sequences declined over time but increased following the frequency of omicron variations. Notably, the optimization and deoptimization of codons in the development of SARS-CoV-2 genomes were shown to be predominantly driven by amino acid alterations, whereas synonymous mutations had minimal impact. We discovered a substantial preference for C>U synonymous changes in the SARS-CoV-2 genomes, and our population genetic simulations suggest mutational pressure might be the primary driving force for this pattern. Through dual fluorescence experiments, we showed that codon optimization through synonymous mutations increases viral protein synthesis and codon deoptimization decreases viral protein synthesis. We also provide experimental evidence that optimizing the codons of the S gene of SARS-CoV-2 variants enhances protein translation in human cells, which has important implications for vaccine design.

The host organisms have evolved multiple defense systems against viruses. The APOBEC family, for example, causes C>U mutations in viral RNA genomes. We found that the C>U mutations are more prevalent in SARS-CoV-2 genomes in human populations than before the outbreak, presumably due to the host shift from non-human animals to humans. Notably, all of the synonymous mutations generated by the C>U substitutions will deoptimize the codons in humans, and this may be a key mechanism human evolved to defend against viruses. On the other hand, the large number of mutations in SARS-CoV-2 caused by the RNA editing systems may supply viruses with an abundance of raw materials for adapting to human host conditions. Thus, the RNA editing mechanisms create a disadvantage that SARS-CoV-2 can exploit. It is probable that SARS-CoV-2 exploits the host’s antiviral system and deoptimizes codon use to better adapt to the host’s immune system.

Instead of developing to utilize more optimized codons in humans, the SARS-CoV-2 exhibited an overall lack of codon optimization. Although intuitively, one expects higher translational efficiency to benefit the virus, it is possible that a high level of translation activity of viral mRNAs will stress the translational machinery of host cells, which does not ultimately benefit the virus. Consistent with this hypothesis, we found the mRNA levels of SARS-CoV-2 tend to be much higher than those of mammalian genes. Specifically, we analyzed the RNA-seq data of Vero E6 cells infected with SARS-CoV-2 and harvested at 4, 6, 12, and 24 hours post-infection (*50*), and the RNA-seq data of Vero cells (ATCC, CCL-81) infected with SARS-CoV-2 and harvested at 24 hours post-infection (*51*). In both datasets, we compared the expression levels (Transcripts Per Kilobase of exon model per Million mapped reads, TPM) of the top 3000 most abundantly expressed cellular genes to those of the SARS-CoV-2 genes. Notably, the TPM values were significantly higher for SARS-CoV-2 genes than cellular genes. In the first dataset (*50*), the median TPM was 61.3, 6,444, 35,049, and 42,061 for the SARS-CoV-2 genes at 4, 6, 12, and 24 hours post-infection, while the median TPM was 68.4, 51.8, 2.42, and 0.2 for the top 3000 most abundantly expressed cellular genes at those time points respectively **(Figure S4)**. Similarly, in the second dataset (*51*), the median TPM was 25,209 for SARS-CoV-2 genes while only 1.136 for the top 3000 most abundantly expressed cellular genes, yielding a ratio of 22,191 **(Figure S4)**. We also retrieved the 23 metagenomic data from the patients. In 20 of these samples, the mRNA levels of SARS-CoV-2 genes were significantly higher than those of the top 3000 most abundantly expressed human genes after correcting multiple tests (**Figure S5**). Moreover, nonstructural protein 1 (nsp1) of SARS-CoV-2 can effectively suppress global translation of host mRNA but not viral mRNAs because of unique feathers of viral 5’ untranslated region (UTR) in translation (*52, 53*). Thus, the SARS-CoV-2 mRNAs are quite abundant in host cells, and it is possible high level of translation activity of viral mRNAs might not be needed for viral replication.

Codon usage affects protein synthesis efficiency, and optimization of codons is vital in vaccine development. The S protein of coronaviruses is the primary target of neutralizing antibodies of host organisms, and several mRNA and protein subunit vaccines of SARS-CoV-2 are intended to target this protein. The different codon usage profiles in SARS-CoV-2 relative to humans emphasized the need to improve the *S* gene’s codon usage for vaccine design against SARS-CoV-2. We demonstrated that codon optimization of SARS-CoV-2 increases protein translation experimentally, and codon optimization substantially increases the levels of S protein expression. Consequently, our findings highlight the necessity of codon usage optimization in mRNA and DNA vaccine formulation.

## Materials and Methods

### SARS-CoV-2 sequence variation and annotation

We downloaded 9,164,789 high-quality SARS-CoV-2 genomes from the Global Initiative on Sharing All Influenza Data (GISAID, https://www.gisaid.org; as of June 18, 2022). We mapped the SARS-CoV-2 genomes to the reference genome (NC_045512) using MAFFT v7.453 (*54*) with the default parameters, and used SnpEff v5.0e(*55*) to annotate the synonymous and nonsynonymous substitutions.

We retrieved the information on the collection date and lineage annotation of the SARS-CoV-2 genomes, and calculated the proportion of VOCs and VOIs in a 14-days window with a one-day step. We also counted the percentage of different nucleotide substitution types in the SARS-CoV-2 genomes collected in each time window. For each synonymous substitution, we calculated the derived allele frequency (DAF) of the synonymous changes in each time window, and used the maximum DAF of a specific change to represent its DAF. We also calculated the DAF of each SNV across all the SARS-CoV-2 genomes.

### Ancestral sequence reconstruction

We downloaded the coding sequences (CDSs) of SARS-CoV-2, RaTG13, BANAL-20-52, Pangolin GX-P5L, and Pangolin GD MP789 from GenBank with the accession number NC_045512, MN996532, MZ937000, MT040335, and MT121216 respectively. We first aligned the protein sequences of each gene using MUSCLE v3.8.31 (*56*) and then reverse-translated them into codon alignments using RevTrans (*57*). We concatenated all the aligned CDS sequences of each genome and reconstructed the phylogenetic tree based on the neighbor-joining algorithm and the Jones-Taylor-Thornton (JTT) model using MEGA X (*58*). The CODEML program in the Phylogenetic Analysis by Maximum Likelihood (PAML) package (*59*) was used to reconstruct the sequences of the most recent common ancestors. The synonymous codon changes from the most recent common ancestor of SARS-CoV-2 and RaTG13 (or BANAL-20-52) to SARS-CoV-2 and RaTG13 (or BANAL-20-52) were defined as pre-outbreak codon changes.

### The calculation of the Codon Adaptation Index (CAI) of SARS-CoV-2

The classic codon adaptive index (CAI) was calculated based on the codon usage profile of the highly expressed genes (*60*). In this study, we calculated expression weighted CAI in humans based on the actual frequency of codon usage in the transcriptomes. We downloaded the gene expression profiles of 54 human tissues from the Genotype-Tissue Expression (GTEx) database Version 8 (https://www.gtexportal.org/). We calculated the median expression value across tissues for each gene and used the median value to represent its expression level. The sequence and annotation for human genome used in this study were downloaded from GENCODE (GRCh38.p13), and the longest transcripts in each gene were retained.

Combining the protein sequences and the mRNA expression levels, we calculated the occurrences of codons in the transcriptome using

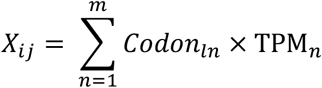

where *X*_*ij*_ represents the occurrence of the *j* th codon of the *i* th amino acid in the transcriptome, *Codon*_*ln*_ represents the occurrence of the *l*th codon in the *n*th gene, TBM_*n*_ denotes the mRNA expression of the *n*th gene, and *m* denotes the number of genes included in the analysis.

Next, we calculated the relative usage of each codon using

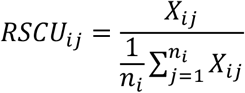

where *n*_*i*_ indicates the number of codon types for the *i* th amino acid. And finally, we calculated the *w*_*ij*_ parameter with

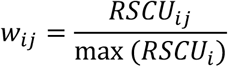

and we further determined the CAI value of a sequence with the geometric mean of the *w*_*ij*_ values for all the codons in that sequence. We defined the codons with *w*_*ij*_ ≥0.9 as optimal and those with *w*_*ij*_ < 0.9 as nonoptimal codons.

We concatenated the coding sequences of SARS-CoV-2 (*ORF1ab, S, ORF3a, E, M, ORF6, ORF7a, ORF7b, ORF8, N*, and *ORF10*) to calculate the expression weighted CAI value. To assess the effects of synonymous mutations on CAI, we replaced the sites with nonsynonymous mutations with the corresponding sites of reference genome in the variant genomes. To obtain the CAI of the native *S* gene of three Omicron lineage (BA.4, BA.5, and BA.2.12.1), we randomly selected one high-quality sequence of each lineage from the GISAID database (BA.4: EPI_ISL_13802998, BA.5: EPI_ISL_13578652 and BA.2.12.1: EPI_ISL_13524674). And the sequences of *S* genes were identified using exonerate v2.4.0 (*61*).

### Population genetics simulations

We used forward simulations to analyze the distribution of derived allele frequencies of different mutational types. Briefly, the simulation begins with 1 randomly generated ancestral sequence with a length (*L*) of 6000 nucleotides, similar to the number of synonymous sites of SARS-CoV-2. The proportions of nucleotide composition in this ancestral sequence were similar to those of the coding sequences of the reference genome of SARS-CoV-2 (A, 0.298582; U, 0.322160; C, 0.183222; G, 0.196036). In the 2nd generation, the ancestral sequence generates 5 random offspring viral sequences following the mutation matrix presented in **Table S2**. Besides considering mutational bias, we assumed different nucleotides to have different mutational rates based on the observations in SARS-CoV-2, with the mutational rates of 2.31×10^−4^, 2.76×10^−4^, 1.284×10^−3^, and 2.58×10^−4^ per site per generation for A, U, C, and G, respectively. On average, the mutational rate used in our simulation was 4.44×10^−4^ per site per generation, which roughly resulted in 2.66 mutations in an offspring sequence. We also assumed the mutational rate of each type of nucleotide to follow a poisson distribution. In the third generation, each of the six viral sequences generates 5 random offspring viral sequences using the same parameter setting. The simulation continues to the next generation with the same parameter. When the population size is above 1,000 in the 5th generation, we randomly selected 1000 parental and offspring sequences in each generation. The simulation stops after 150 generations. The whole simulation process was replicated 100 times, and the distribution of derived allele frequency of each mutational type is summarized based on the 100 replicates of simulations.

### Dual-luciferase reporter assay

Dual-luciferase assay (psiCHECK-2 vector, Promega) was selected to analyze the effects of the viral mutation on protein synthesis. The primers psi_backbone_F and psi_backbone_R were used to generate PCR products (the vector’s backbone) with appending overhangs. Following gel extraction, template plasmids were digested using *DpnI* (NEB). The wildtype and mutated viral sequence were constructed using overlap PCR, and assembled with the backbone using the NEBuilder HiFi DNA Assembly Cloning Kit (E5520S, NEB). All primers used in this study were listed in Supplementary **Table S3**.

HEK293FT cells were purchased from the Cell Bank of the Chinese Academy of Sciences, and were maintained in Dulbecco’s Modified Eagle medium (DMEM, Gibco) with 10% FBS (Gibco) and 1% penicillin-streptomycin at 37 °C in a humid incubator with 5% CO2 in the air. Plasmids were extracted using the QIAGEN Miniprep kit (27106, QIAGEN) as per manufacturers’ instructions. Constructed vectors were transfected into HEK293FT cells using Lipofectamine 3000 Transfection Reagent (L3000015, Thermo Fisher Scientific). Cells were cultivated for 32h after transfection, and then a dual-luciferase reporter assay system (Promega) was used to detect Renilla levels of wildtype and mutant plasmids and normalized to Firefly luciferase as an internal control (five biological repetitions per plasmid).

### Western blot analysis of S protein expression

The individual BA.2.12.1 and BA.4/5 S protein encoding sequences with various CAIs were synthesized by GeneScript (Nanjing, China) and cloned into pcDNA 3.1 plasmid between *Hind* III and *Xba* I sites. HEK293T cells were obtained from ATCC and maintained in DMEM with 10% FBS and 1% penicillin-streptomycin at 37 °C and 5% CO_2_. About 2×10^6^ cells of HEK293T cells were transfected with 2 ug of plasmids encoding S proteins using polyetherimide (PEI; Sigma). After 16 hours of incubation, cells were fed with fresh medium. The next day, cells were lyzed with cell lysate buffer (20 mmol/L Tris-HCl pH 7.5, 150 mmol/L NaCl, 1 mmol/L EDTA, 0.1% sodium dodecyl sulfate (SDS), 1% NP40, 1x protease inhibitor cocktail (Bimake, Houston, USA), and centrifuged at 12,000 g for 10min to remove nuclei. The cell lysates were boiled and separated in 10% SDS-polyacrylamide gel electrophoresis (PAGE) gel. After being transferred to nitrocellulose filter membranes (GVS, Sanford, USA), the samples were detected using polyclonal rabbit anti-SARS-CoV-2 S antibodies at a dilution of 1:2,000.

### Quantification of viral and cellular gene expression

We retrieved the RNA-seq data of Vero E6 cells infected with SARS-CoV-2 at a multiplicity of infection (MOI) of 0.1 and harvested at 4, 6, 12, and 24 hours post-infection (*62*), and the RNA-seq data of Vero cells (ATCC, CCL-81) infected SARS-CoV-2 at an MOI of 0.05 and harvested at 24 hours post-infection (51). We calculated the gene expression of SARS-CoV-2 and the hosts using the Kallisto program (*63*), with the longest transcript of *Chlorocebus sabaeus* (ChlSab1.1) and the CDS of SARS-CoV-2 (NC_045512) as the reference transcriptomes. We also retrieved the 36 metagenomic data of the patients from NCBI short read archive (SRA), and kept 23 libraries with gene expression information for all the SARS-CoV-2 genes. The expression levels of SARS-CoV-2 and human genes were also calculated by the Kallisto program (*63*).

## Acknowledgments

We thank the researchers who generated and shared the sequencing data from GISAID (https://www.gisaid.org/) on which this research is based. This work was supported by the National Key Research and Development Projects of the Ministry of Science and Technology of China (2021YFC2301300, 2021YFC0863300, 2020YFA0707600, 2020YFC0847000), SLS-Qidong Innovation Fund, and Chinese Academy of Medical Sciences Innovation Fund for Medical Sciences (2021-12M-1-038).

## Supplementary Tables and Figures

**Table S1.**
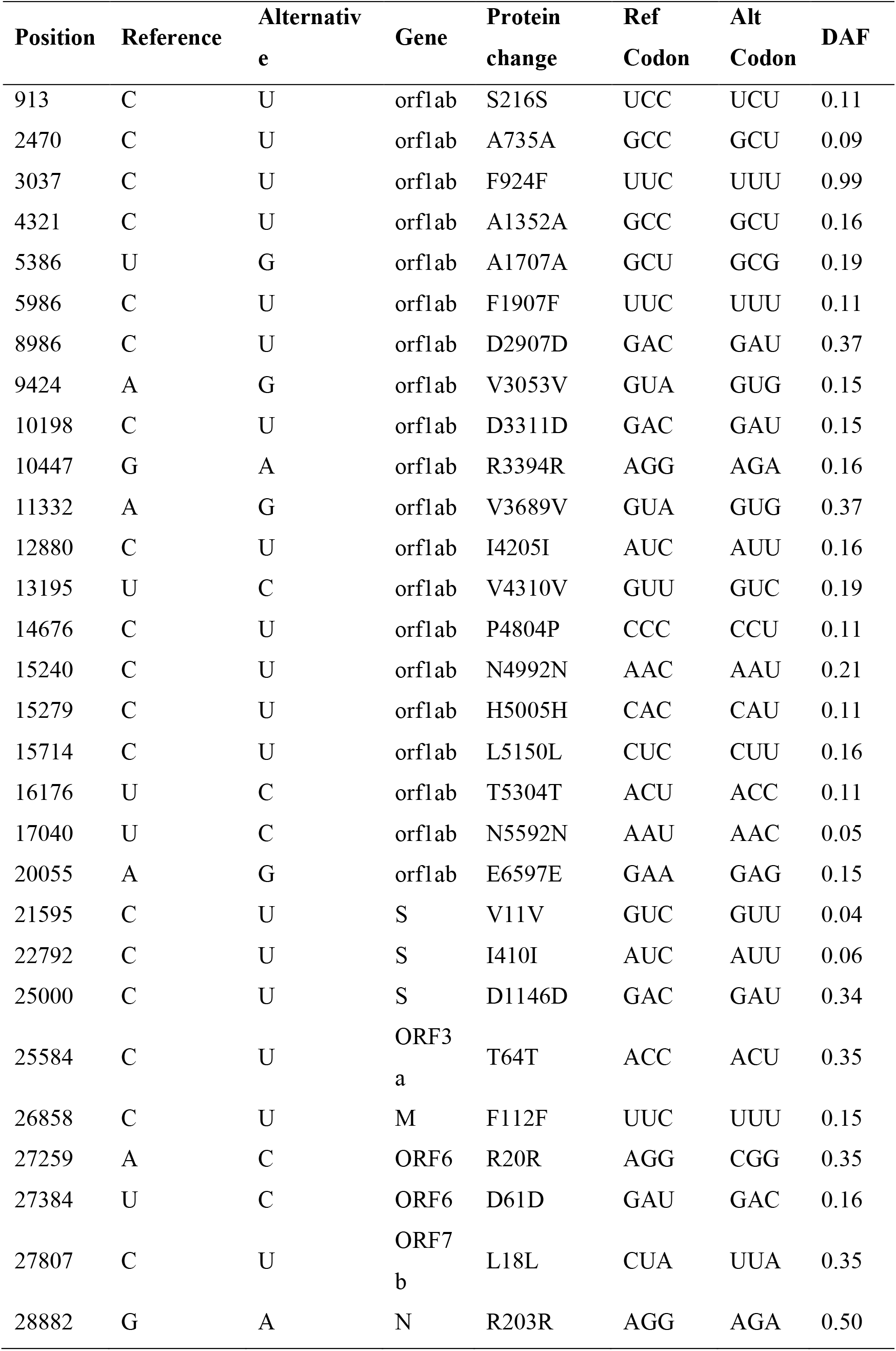
The synonymous mutations with higher DAF values.

| Position | Reference | Alternative | Gene | Protein change | Ref Codon | Alt Codon | DAF |
| --- | --- | --- | --- | --- | --- | --- | --- |
| 913 | C | U | orf1ab | S216S | UCC | UCU | 0.11 |
| 2470 | C | U | orf1ab | A735A | GCC | GCU | 0.09 |
| 3037 | C | U | orf1ab | F924F | UUC | UUU | 0.99 |
| 4321 | C | U | orf1ab | A1352A | GCC | GCU | 0.16 |
| 5386 | U | G | orf1ab | A1707A | GCU | GCG | 0.19 |
| 5986 | C | U | orf1ab | F1907F | UUC | UUU | 0.11 |
| 8986 | C | U | orf1ab | D2907D | GAC | GAU | 0.37 |
| 9424 | A | G | orf1ab | V3053V | GUA | GUG | 0.15 |
| 10198 | C | U | orf1ab | D3311D | GAC | GAU | 0.15 |
| 10447 | G | A | orf1ab | R3394R | AGG | AGA | 0.16 |
| 11332 | A | G | orf1ab | V3689V | GUA | GUG | 0.37 |
| 12880 | C | U | orf1ab | I4205I | AUC | AUU | 0.16 |
| 13195 | U | C | orf1ab | V4310V | GUU | GUC | 0.19 |
| 14676 | C | U | orf1ab | P4804P | CCC | CCU | 0.11 |
| 15240 | C | U | orf1ab | N4992N | AAC | AAU | 0.21 |
| 15279 | C | U | orf1ab | H5005H | CAC | CAU | 0.11 |
| 15714 | C | U | orf1ab | L5150L | CUC | CUU | 0.16 |
| 16176 | U | C | orf1ab | T5304T | ACU | ACC | 0.11 |
| 17040 | U | C | orf1ab | N5592N | AAU | AAC | 0.05 |
| 20055 | A | G | orf1ab | E6597E | GAA | GAG | 0.15 |
| 21595 | C | U | S | V11V | GUC | GUU | 0.04 |
| 22792 | C | U | S | I410I | AUC | AUU | 0.06 |
| 25000 | C | U | S | D1146D | GAC | GAU | 0.34 |
| 25584 | C | U | ORF3<br>a | T64T | ACC | ACU | 0.35 |
| 26858 | C | U | M | F112F | UUC | UUU | 0.15 |
| 27259 | A | C | ORF6 | R20R | AGG | CGG | 0.35 |
| 27384 | U | C | ORF6 | D61D | GAU | GAC | 0.16 |
| 27807 | C | U | ORF7<br>b | L18L | CUA | UUA | 0.35 |
| 28882 | G | A | N | R203R | AGG | AGA | 0.50 |

**Table S2.** The mutational matrix used for population genetics simulation.

|  |  |  |  |  |  |
| --- | --- | --- | --- | --- | --- |
|  |  | **Ancestral allele** |  |  |  |
|  |  | **A** | **U** | **C** | **G** |
| **Derived allele** | **A** | 0.999769 | 0.000012 | 0.000010 | 0.000180 |
|  | **U** | 0.000015 | 0.999724 | 0.001259 | 0.000062 |
|  | **C** | 0.000035 | 0.000243 | 0.998716 | 0.000016 |
|  | **G** | 0.000181 | 0.000021 | 0.000015 | 0.999742 |

**Table S3 is an excel file that contains the primer sequences for PCR**.

**Figure S1.**
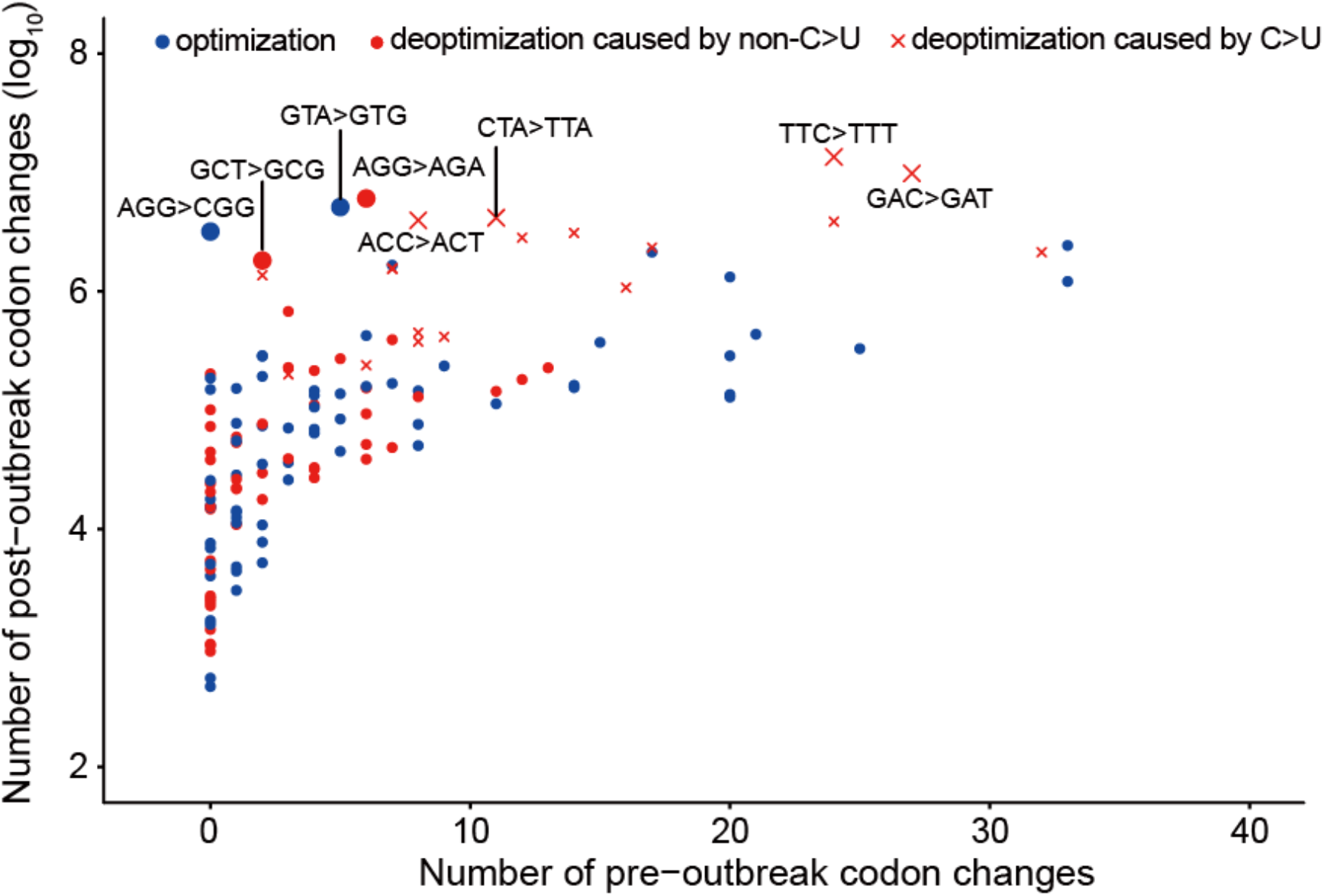
The post-outbreak (*y*-axis) over pre-outbreak (*x*-axis) synonymous codon change. The pre-outbreak codon changes were inferred based on the synonymous codon changes from the most recent common ancestor of BANAL-20-52 and SARS-CoV-2 to them in the pre-outbreak evolution history.

**Figure S2.**
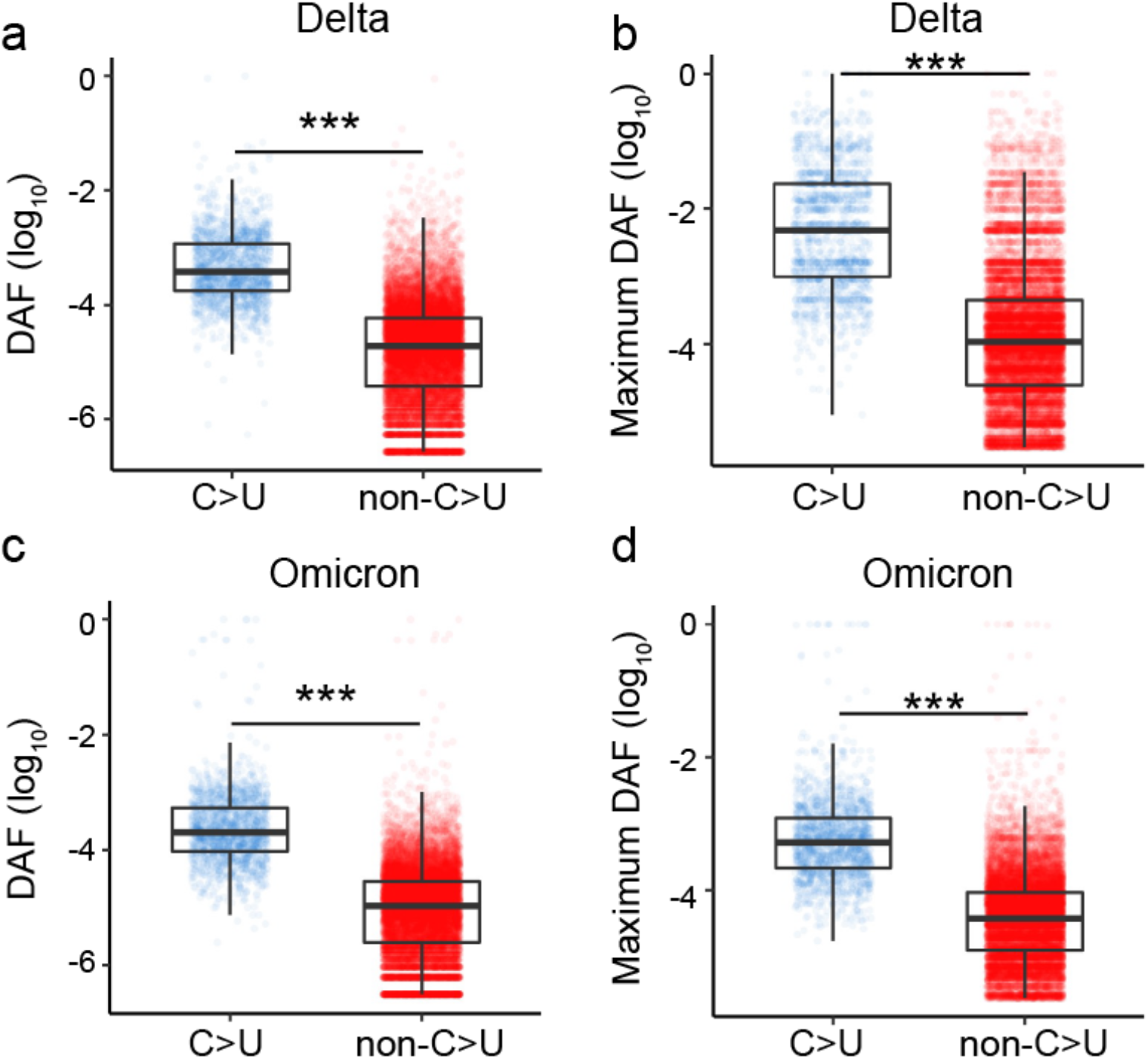
The DAFs of the C>U synonymous changes are significantly higher than those of the non-C>U synonymous changes in the Delta (a, b) or Omicron variants (c, d). (a, c) All the mutations were considered in the mutation frequency calculation. (b, d) The maximum DAF in a time window was used to represent the DAF of a synonymous mutation.

**Figure S3.**
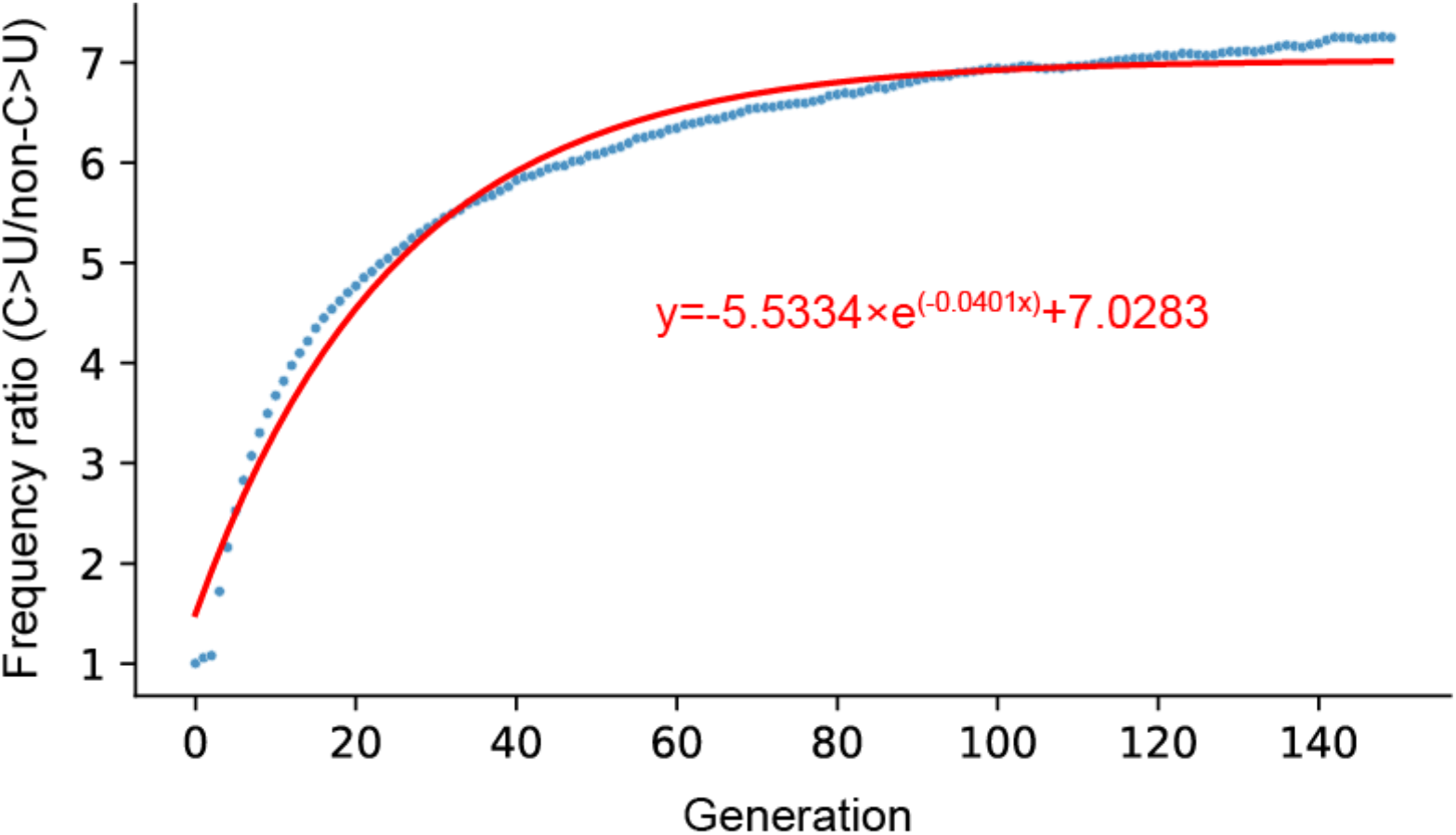
The ratio of DAF corresponding to C>U mutations and non-C>U mutations during the simulation. The blue dots are the ratio of the frequencies of the two types of mutations in each generation. The red line is the fitted line.

**Figure S4.**
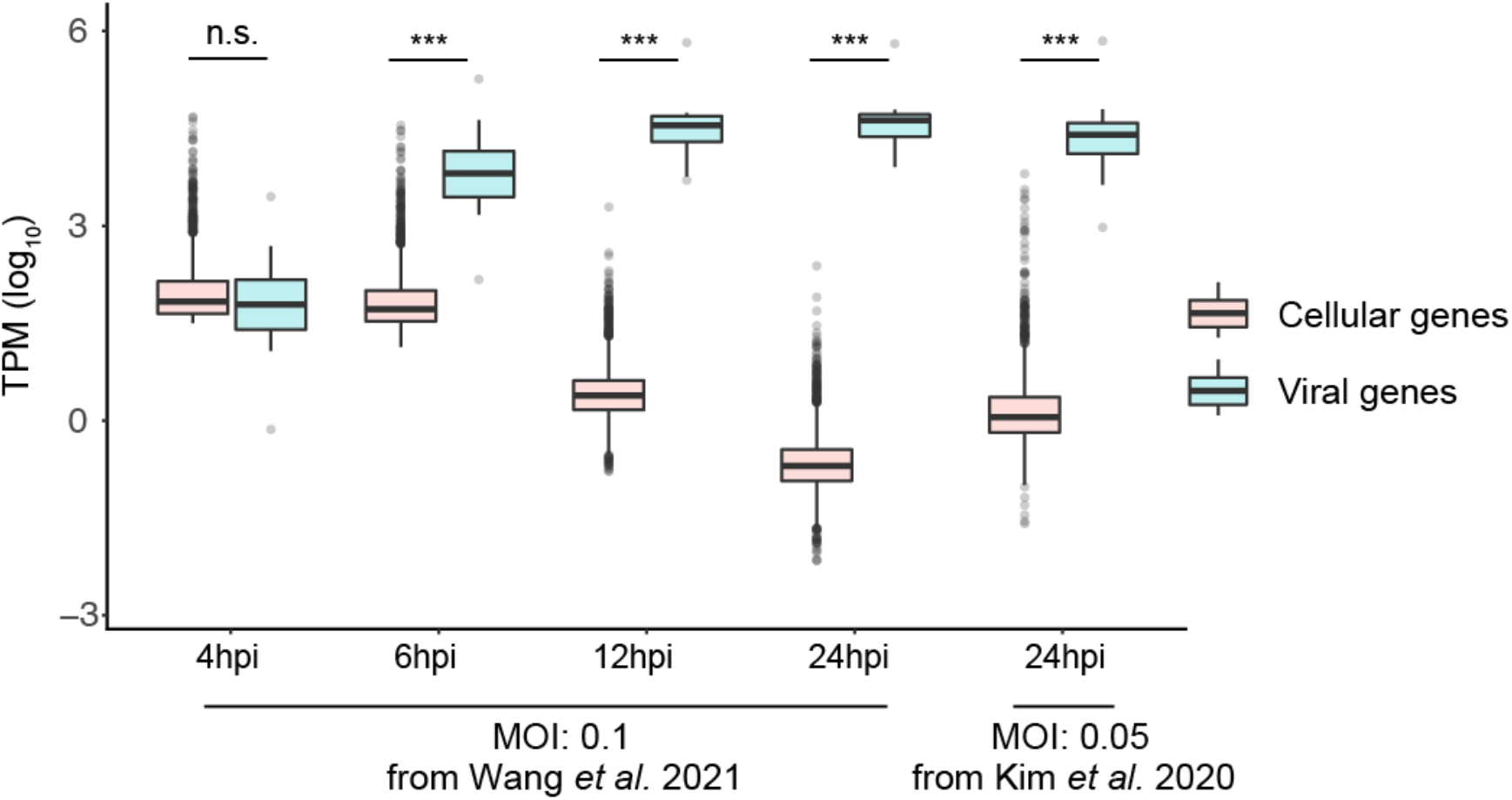
The gene expression levels of Vero cells infected with SARS-CoV-2 and harvested at different time points. For the Vero cells, only the top 3000 highly expressed genes were considered in the analysis. The initial multiplicity of infection (MOI) and the cultivation time (hours post-infection, hpi) were labeled on the *x*-axis. ***, *P* < 0.001.

**Figure S5.**
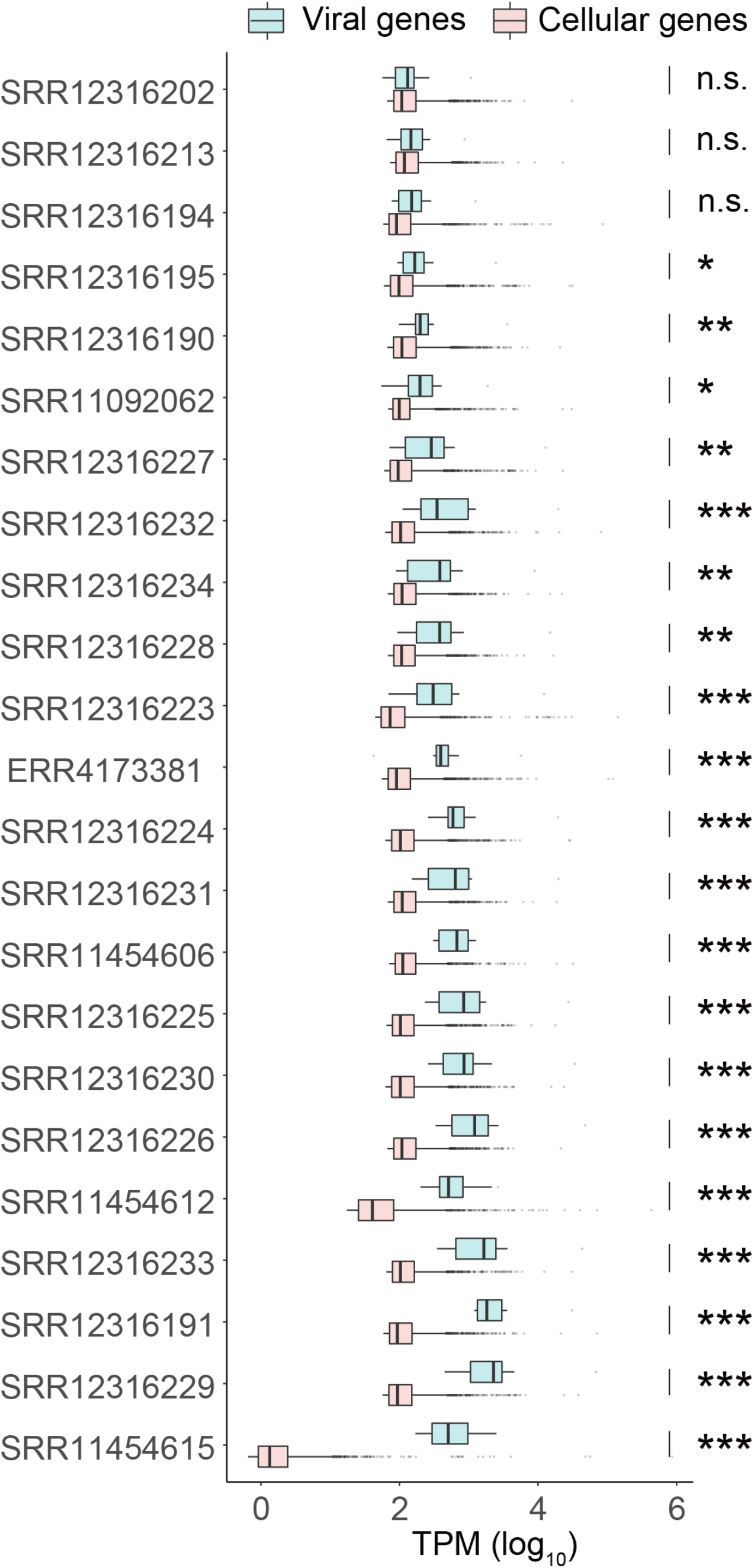
The gene expression levels of SARS-CoV-2 and human patients detected from metagenomic data. ***, *P* < 0.001.

